# Connectivity patterns predictive of cognition, but not affect, reflect a segregated intrinsic network architecture

**DOI:** 10.64898/2026.02.10.704998

**Authors:** Achille Gillig, Gaël Jobard, Sandrine Cremona, Marc Joliot

## Abstract

The brain’s intrinsic organization into resting-state networks has long been suggested to be fundamental for the offline support of mental processes. Extensive task-based evidence support the relevance of the crosstalk between network segregation, supporting systems specialization, and network integration, allowing to flexibly implement complex behavior. However, only scarce evidence focusing on few behavioral measures directly link changes in these network properties at rest with interindividual differences in behavior.

In this work, using a comprehensive set of behavioral measures together with resting-state functional magnetic resonance imaging from the human connectome project, we assessed whether connectivity patterns predictive of behavior reflected segregation or integration based on GINNA, a 33 resting-state-networks atlas with cognitive characterization.

We found that connectivity relevant for behavior organizes into 3 main latent dimensions, summarizing *Cognition*, *Positive Affect* and *Negative Affect*. Crucially, we found that connectivity predictive of *Cognition*, but not *Affect*, was associated with global network segregation and reduced network integration. We further reveal differential resting-state-networks involvements, with *Cognition* associated with the segregation of higher-level resting-state-networks, and the integration of lower-level, visual networks.

All in all, the present results suggest that cognition may rest upon a segregated, modular intrinsic brain architecture.

**Author summary:** How psychological processes are maintained when the brain is not engaged in an explicit task remains unclear. Resting-state networks offer a candidate intrinsic architecture, but whether interindividual differences in diverse behavioral domains relate to this architecture in similar ways is unknown. We combined Human Connectome Project resting-state fMRI with 58 behavioral measures to identify functional connectivity patterns predictive of behavior and examine how they relate to network segregation and integration. These patterns organized into three latent dimensions summarizing Cognition, Positive Affect, and Negative Affect. Cognition-related patterns aligned with a segregated intrinsic architecture, showing stronger within-network and weaker between-network connectivity, particularly among higher-order resting-state networks, with complementary integration of visual networks. Affective patterns showed less consistent network signatures, suggesting a more distributed or context-dependent intrinsic organization.

## 1 Introduction

The intrinsic organization of the brain into networks, and how it relates to mental processes, has attracted much attention. Using resting-state functional magnetic resonance imaging (rs-fMRI), it was shown that the brain is intrinsically organized into networks, i.e., groups of regions with tightly coupled spontaneous blood oxygen-level dependent (BOLD) signal fluctuations (Biswal et al., 1995; Fox et al., 2005; Fox & Raichle, 2007). These so-called resting-state networks (RSNs), ever since their first observations (Biswal et al., 1995), have been suggested to hold a functional relevance with respect to cognitive processes and behavior, largely because their topology follows the boundaries of cognitive systems (Seitzman et al., 2019; Uddin et al., 2023). This strong assumption is nowadays reflected in the tendency for researchers to refer to RSNs by their putative cognitive functions, rather than an anatomically grounded taxonomy (Uddin et al., 2019, 2023).

Support for the notion that RSNs represent fundamental cognitive modules was initially gathered from comparisons with tasks, with a first evidence of a similar network architecture between rest and task (Smith et al., 2009). Additional evidence showed that task network architecture is shaped by RSN architecture (Cole et al., 2014, 2016) and that RSNs display modularity with respect to their task engagement (Bertolero et al., 2015; Yeo et al., 2015). Motivated by these evidence, a 33 resting-state networks atlas grounded in meta-analytical cognitive characterization was recently proposed, summarizing potential associations with cognition in light of the neuroimaging literature (Gillig et al., 2025). Yet, while informative, spatial comparison between RSNs and task activation maps provide evidence only of a coarse, associative, but not an exploratory, relationship (Mill et al., 2017).

More direct evidence come from the study of resting-state functional connectivity (FC), measuring the strength of the functional coupling between pairs of regions or networks (typically, measured as the correlation between resting-state timeseries). FC is now widely used to predict a wide range of behavior and latent cognitive processes, including, but not limited to, intelligence (Thiele et al., 2024), general cognition (Ooi et al., 2022), working memory (Avery et al., 2020), or personality (Nostro et al., 2018), indicating that individual differences are reflected as long-term changes in the brain functional organization (Stevens & Spreng, 2014; Wu et al., 2023).

While studies linking variations in FC to behavior are now many, those that probe the relevance of the intrinsic RSN architecture are scarce. Switching from a resting to a cognitive state implies large-scale network reconfigurations, *i.e.*, regions change their network affiliations to flexibly adapt to ongoing demand (Salehi et al., 2020). However, it has been shown that decreased task-related network reconfiguration was associated with better task performance across language, reasoning and working memory (Schultz & Cole, 2016), suggesting that RSNs provide a scaffold for efficient active cognitive processing.

Taken together, these results outline a framework in which resting-state FC may index the offline maintenance of behavior through mechanisms of Hebbian plasticity, where repeated task-related co-activation strengthens intrinsic functional couplings. In this framework, RSNs scaffold cognition because behaviorally relevant couplings increasingly align with RSNs delineations, therefore constraining and facilitating subsequent task-evoked activations.

To probe the relevance of the intrinsic RSN organization with respect to behavior, network theory has proven to offer interesting tools. Of note, the brain constantly alternates between integrated (increased cross-network communication) and segregated (increased network independence) states, both at rest and during tasks (Capouskova et al., 2023; Cohen & D’Esposito, 2016; Shine et al., 2016; Shine & Poldrack, 2018; Wang et al., 2021), with segregation linked to the maintenance of a modular, specialized information processing architecture (Sporns & Betzel, 2016), while integration allows cross-systems communication to meet complex task demands (Bassett et al., 2011; Bertolero et al., 2015, 2018; Capouskova et al., 2023; Cohen & D’Esposito, 2016; Shine et al., 2016).

Most evidence linking network segregation and integration to cognitive performance comes from task-based studies. During active task performance, greater integration has been associated with faster and more accurate performance under high cognitive demand (Shine et al., 2016). Consistently, a relationship with task complexity was established: higher efficiency correlated with increased network integration during a working memory task, whereas it was associated with greater segregation for a simpler motor task (Cohen & D’Esposito, 2016). Similarly, in the context of stimulus-response (S-R) learning, it was found that initial, goal-directed learning involved increased integration, while later, automated S-R associations were associated with increased segregation (X. Wang et al., 2024). Altogether, these task-based findings highlight the importance of the balance between network segregation to promote, local, specialized within-network processing for overlearned processes, and integration to flexibly adapt to changing demands (Cohen & D’Esposito, 2016; Shine & Poldrack, 2018).

While a clear picture starts to emerge when it comes to task-related cognitive processing, highlighting the relevance of both segregation and integration mechanisms, evidence on how their interplay promotes the offline, at-rest maintenance of behavior remains scarce. While the resting state was found to be characterized by a balance between segregated and integrated states (Wang et al., 2021), individual differences in segregation and integration were also linked to cognitive abilities. Stronger integration was associated with better general cognitive ability, while crystallized intelligence and processing speed involved stronger segregation, and a tendency towards balance supported better memory (Wang et al., 2021). As this suggests opposed effects between the maintenance of global versus narrower processes, questions remain regarding a wider range of cognitive, but also socio-emotional and personality functioning, and to what extent their supporting mechanisms are shared or unique across this diverse range of behavioral domains. In addition, it remains to be determined whether these mechanisms differ across RSNs as a function of the behavioral domain involved.

In this study, using data from the Human Connectome Project, we investigated whether individual differences in multiple domains of behavior were predicted by intrinsic FC patterns, and whether these patterns related to specific integration/segregation balance across different RSNs.

First, we performed FC prediction of 58 behavioral measures spanning cognitive, socio-emotional, and personality domains. The Haufe method (Haufe et al., 2014) was used to gain insights into the relevance of region-level FC. We next performed exploratory factor analysis to explore the latent FC structure associated with behavior. Last, a reliability analysis was conducted both at the global and RSN level to identify whether latent FC factors were associated with reliable changes in network segregation and integration across components of a cognitively characterized RSN atlas.

## 2 Material and Methods

### 2.1 Participants

We analyzed data from the HCP Young Adult Sample S1200 (Smith et al., 2013) including 1,200 subjects (age: 22–37 years; 656 female). Study procedures were approved by the Washington University Institutional Review Board, and informed consent, in accordance with the Declaration of Helsinki, was obtained from all participants. Subjects with missing behavioral data for any of the 58 selected behavioral variables (Liégeois et al., 2019) (Supplementary Table 1) were excluded, resulting in 1,022 participants with complete behavioral assessment. After fMRI data quality checks (see Resting-state fMRI acquisition and preprocessing) 779 participants were included for subsequent analyses (22–37 years, mean age = 28.67 years; 401 female), representing a well-power sample size for modeling prediction (Ooi et al., 2025).

### 2.2 Behavioral measures

We selected 58 commonly used behavioral measures (Liégeois et al., 2019) spanning cognitive, socio-emotional, and personality domains (Supplementary Table 1) mainly originating from the NIH toolbox, and NEO-FFI for personality (Barch et al., 2013).

### 2.3 Resting-state fMRI acquisition and preprocessing

The four runs (two sessions, each with LR and RL acquisition) of the resting-state fMRI data present in the HCP dataset were used for analyses. Details regarding the rs-fMRI acquisition are available in Smith et al. (Smith et al., 2013). Briefly, fMRI data were acquired on a 3-T Siemens Skyra with a 32-channel head coil and a gradient-echo echo-planar imaging (EPI) sequence (repetition time: TR = 720 ms, echo time: TE = 33.1 ms, flip angle = 52°, 2-mm isotropic voxel resolution, multiband factor = 8). The minimally preprocessed rs-fMRI data was recovered and each of the two runs (LR and RL acquisition) corresponding to each session (REST1 and REST2) were separately concatenated and subjected to noise regression with 24 head motion parameters, eight mean signals from white matter and cerebrospinal fluid, and four grey matter signals. Global signal regression was used based on evidence that it improves performances for functional connectivity prediction of behavior (Li et al., 2019). In-scanner head motion was measured by framewise displacement (FD), and, similarly to Thiele et al. (2024), subjects were only included if mean FD < 0.2 mm, proportion of spikes (FD > 0.25 mm) < 20%, and no spikes were above 5 mm (Parkes et al., 2018). Frames with above-threshold FD (> 0.2 mm) were censored and replaced by an interpolation of the two neighboring frames. Fully preprocessed rs-fMRI data were further parcellated into 448 regions of interest (Cremona et al., 2025) (Fig. 1), resulting in 448 timeseries averaged by parcels.

**Fig. 1.**
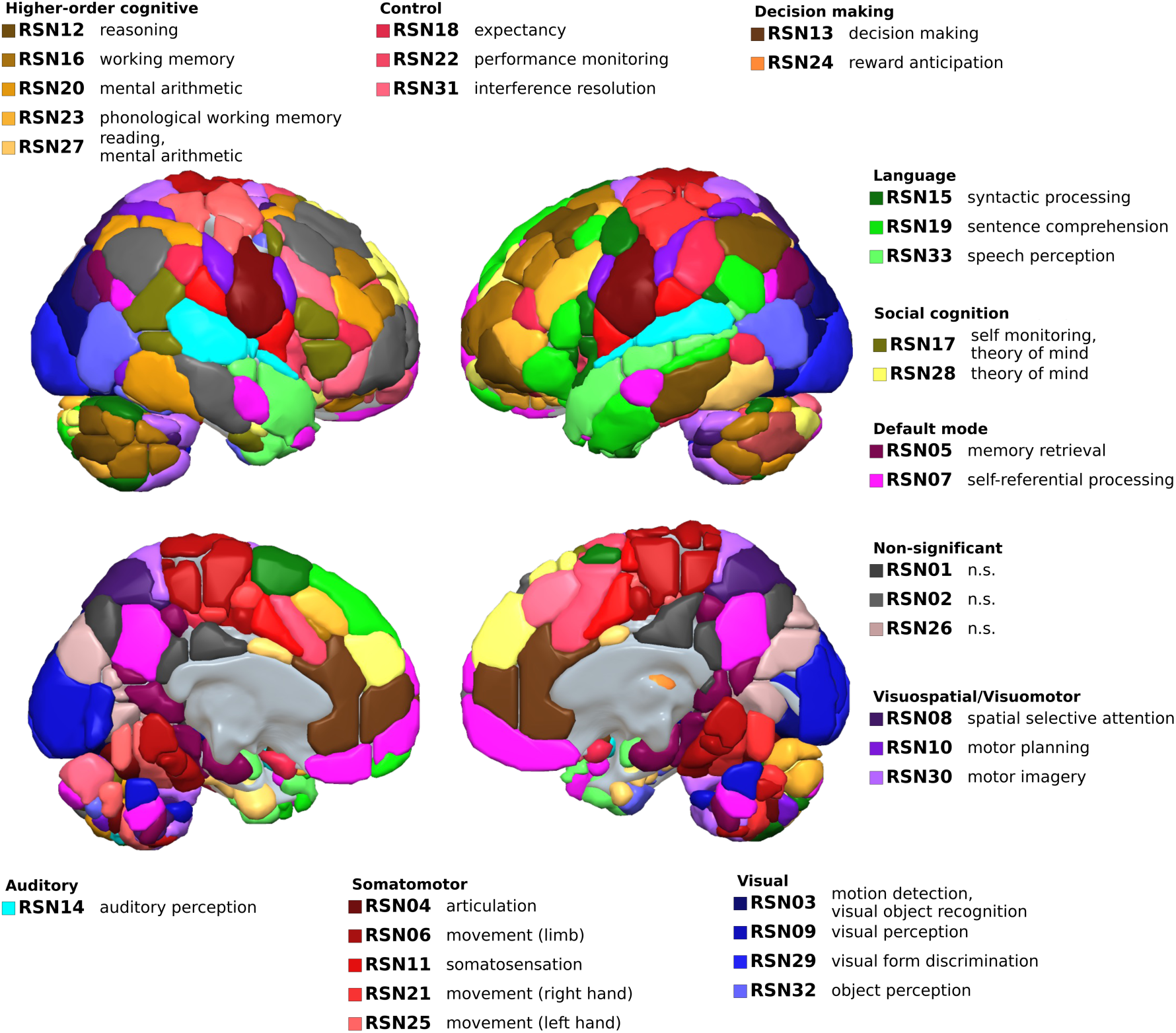
Parcellation and resting-state-networks atlas. A 448 parcellation atlas was employed (Cremona et al., 2025), with each parcel mapped to one of 33 cognitively characterized GINNA RSNs (Gillig et al., 2025) or 1 brainstem network (not shown for visualization purposes). RSNs are categorized according to the cognitive domains defined in Gillig et al. (2025). Abbreviations: n.s., non-significant.

### 2.4 Resting-state networks definition

To define resting-state-networks (RSNs), the Groupe d’Imagerie Fonctionnelle Network Atlas (GINNA) was used (Gillig et al., 2025) (Fig. 1). The 448 parcels of the region-level parcellation atlas (Cremona et al., 2025) are assigned to one of the 33 networks (32 cortical, 1 subcortical) included in GINNA (Gillig et al., 2025), in addition to 1 brainstem network. Similarly to Cremona et al. (2025), we used a version of this atlas without overlap (1 region belongs to 1 RSN), such that FC measurements are not cross-dependent between RSNs. The choice of this specific network atlas was motivated by i) its previous cognitive characterization relying on neurosynth-based meta-analytic decoding, allowing to put into perspective RSNs relationship with behavior at rest in light of their task involvement; ii) its higher granularity, potentially providing finer associations between behavior and RSNs.

### 2.5 Functional connectivity

Functional connectivity (FC) was computed for each individual participants and each acquisition session using Pearson correlation as a metric, followed by Fisher r-to-z transform, resulting in 448×448 FC matrices for each individual and each of the two sessions of rs-fMRI acquisition. Last, FC estimates for the two sessions were averaged to produce a single functional connectivity estimate per subject. While it is more common to concatenate all rs-fMRI data prior to computing functional connectivity, this method was shown to be comparable to averaging out after computing functional connectivity for each individual scans (Cho et al., 2021).

### 2.6 Functional connectivity prediction of behavior

To predict each of the 58 selected behavioral measures, we employed a nested cross-validation procedure with linear kernel ridge regression (Fig. 2A). This model was chosen as it has been demonstrated to achieve similar performance as deep learning approaches (He et al., 2020) while reducing by several orders of magnitude the computational cost.

**Fig. 2.**
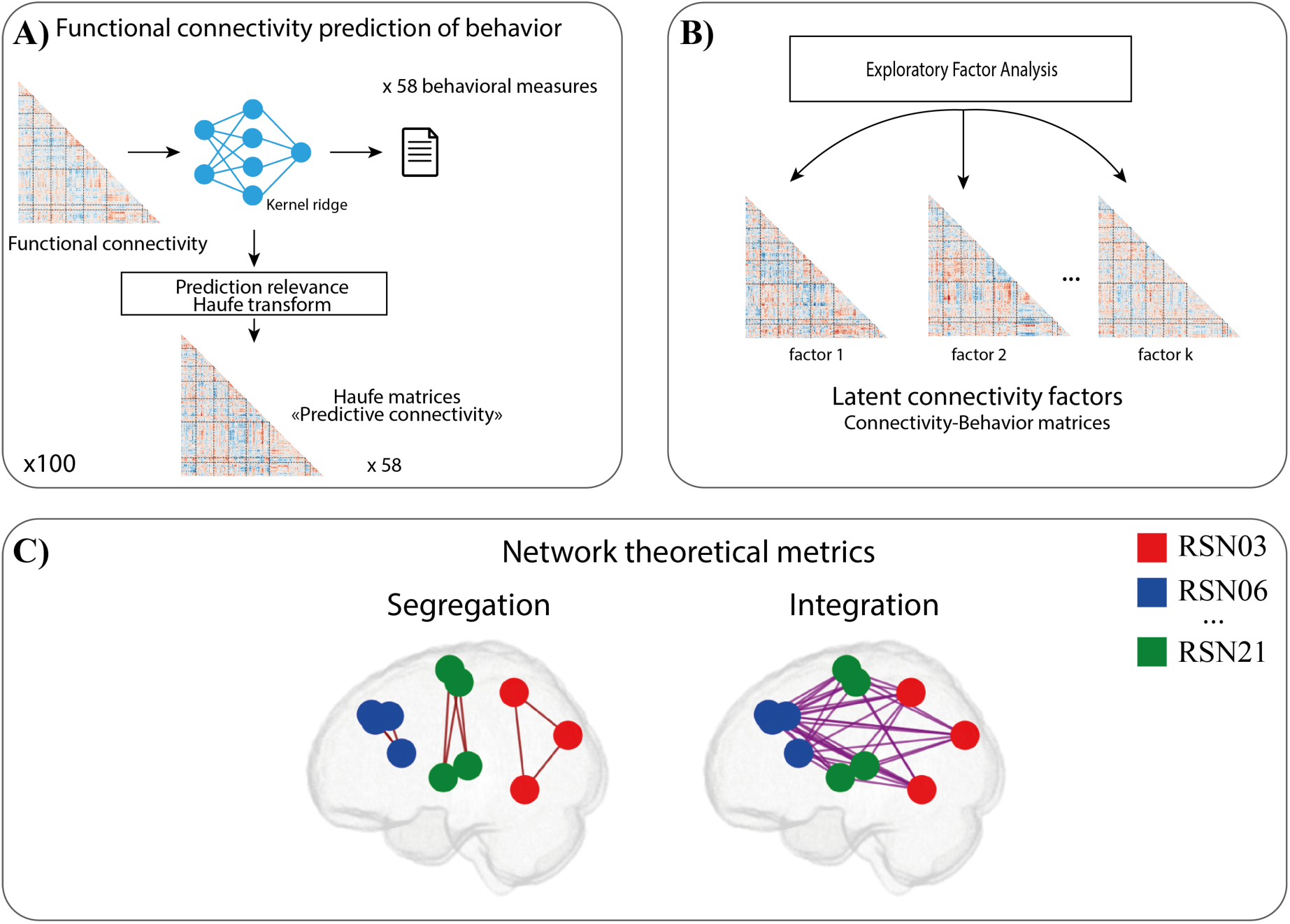
Analysis pipeline. A) Functional connectivity prediction of behavior. 58 behavioral measures were predicted from functional connectivity, with 100 repetitions with varying split-half sampling. The Haufe model inversion method5 was used to retrieve predictive connectivity matrices associated with each behavior. **B) Latent connectivity-behavior factors.** Exploratory factor analysis was used to extract latent predictive connectivity factors. **C) Network theoretical metrics.** Measures of network segregation and integration were computed for each latent connectivity factors: network segregation was defined as the average connectivity between all pairs of regions belonging to the target network, while network integration was defined as the average connectivity between regions belonging to the target network and regions belonging to all other networks.

We used a cross-validation scheme analogous to the one described by Tian & Zalesky (2021), namely, a repeated split-half with nested 10-folds cross-validation procedure. First, participants were split in half. The first half was used as training set, and split into 10 folds for cross-validated hyperparameter optimization (regularization parameter alpha, 5 values on a logscale ranging from −3 to 3). The optimal model was further retrained on the whole training set and used to predict behavioral scores from functional connectivity of the second half (test) set. The same procedure was repeated using the second half as training and the first half as testing, such that each split-half iteration resulted in two estimates of performance.

This procedure was repeated 100 times by varying the split-half sampling (Tian & Zalesky, 2021) in order to obtain stable performance estimates (Varoquaux et al., 2017). The final performance estimation was obtained by averaging across the 100 repetitions. Control over confounding variables was performed following a procedure of regression over the output of the prediction (predicted y) (Dinga et al., 2020).

### 2.7 Prediction performance significance

Performance for predicting participants’ behavioral scores from FC was measured as the standardized root mean squared error (sRMSE) (i.e, RMSE divided by variance) computed after accounting for confounds (Dinga et al., 2020).

To assess whether each behavioral measure could be predicted from FC better than chance, we compared the distribution of prediction errors to a distribution of null prediction errors generated by permutations. FC matrices were randomly permuted and the whole nested cross-validation procedure was repeated. Importantly, confounds were not permuted, so as to preserve the relationship between confounds and predicted variables (Dinga et al., 2020). Similarly to the empirical predictions, the procedure was iterated 100 times, producing a distribution of prediction performance that would be obtained if there was no association between FC and behavior. Due to the half-split cross-validation scheme, each iteration produced two estimates, resulting in a null distribution of n = 200. For each latent factor, the observed performance estimates were compared to the null distributions using a paired t-test (Tian & Zalesky, 2021), followed by Benjamini-Hochberg false discovery rate correction for multiple comparisons (Benjamini & Hochberg, 1995) (q < 0.05). All non-significant behavioral measures were excluded from downstream analyses.

### 2.8 Prediction relevance interpretability features

For each of the 100 resampling of the half-split procedure, the Haufe model inversion method (Haufe et al., 2014) was used to reveal the contribution of each feature of the whole-brain functional connectome (Fig. 2A). We used Haufe-transformed weights rather than connectome-based predictive mapping (CPM) (Shen et al., 2017) because our aim was to obtain connectivity patterns suitable for downstream latent-factor and network-level analyses. Although CPM is a well-established framework for connectome-based prediction, Haufe-transformed weights can be combined with kernel ridge regression and have been reported to provide reliable feature-level estimates for interpretation (Tian & Zalesky, 2021).

Haufe coefficients were obtained as the following (Haufe et al, 2014):

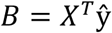

With *X* ∈ *R*^n,m^ the FC matrices of n training participants of length m edges, ŷ ∈ *R*^n^ the predicted behavioral measure for the training set.

This resulted, for each edge, in a coefficient interpretable in a similar manner as FC (Haufe et al., 2014) where a positive coefficient implies a positive relationship between FC and behavior across individuals, and a negative coefficient the inverse relationship. Haufe matrices were further standardized by dividing by the variance of ŷ to allow for comparisons between behavioral measures (Chen et al., 2022).

### 2.9 Latent connectivity-behavior

To reveal the latent organization of FC associated with behavior, an exploratory factor analysis (EFA) with varimax rotation was applied (Fig. 2B). Haufe patterns from the 100 prediction repetitions (n = 200, due to the half-split scheme) were first split in half. The first half (n = 100) was used to compute EFA, while the obtained loadings were thresholded to retain only strongest loadings (|*λ*| > 0.3) (Peterson, 2000) and subsequently applied to the second half to reconstruct latent connectivity-behavior (CB) matrices as follows:

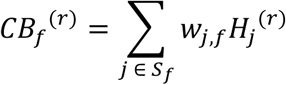

where *H^(r)^_j_* denotes the Haufe matrix for behavioral measure j and repetition *r*, *S_f_* is the set of behavioral measures with retained loadings on factor *f,* and *w_j,f_* is the normalized loading of behavioral measure *j* on factor *f*. The weights were obtained by normalizing the thresholded loadings as follows:

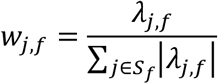

Such that

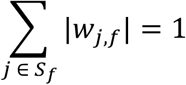

while preserving the sign of each loading.

The optimal number of factors was estimated using parallel analysis (Horn, 1965) with 100 iterations. Consistent with Costello & Osborne (2005) recommendations on factor retention, factors with minimal contribution to explained variance were excluded to favor a parsimonious solution.

### 2.10 Resting-state networks segregation and integration

For each of the k latent CB factors, we investigated network segregation and network integration, both at a global level and at a RSN level (Fig. 2C). Importantly, segregation and integration are not measured directly from participants’ FC matrices, but are derived from predictive connectivity (Haufe) coefficients, and therefore reflect how connectivity patterns associated with behavior are organized. Network integration was operationalized as the average CB patterns between regions belonging to the target RSN and all other regions, while network segregation was operationalized as the average CB patterns of all regions within the target RSN (Fig. 2C). Positive values for integration or segregation indicated a positive association with higher values of the latent factor, while negative values were predictive of lower values of the factor. Global (pooled) estimates of network segregation and integration were obtained by averaging estimates across RSNs. Network metrics significance was assessed post-hoc, based on their reliability across prediction repetitions (95% confidence interval).

We operationalized strong evidence for a segregated CB pattern as the joint presence of positive within-network CB weights (segregation) and negative between-network CB weights (integration), indicating that higher behavioral scores are associated with stronger within-network connectivity and weaker between-network connectivity. This definition was preferred because it preserves the respective contributions of intra- and inter-network connectivity and avoids additional thresholding and sign-handling choices when applied to signed FC-derived matrices. As a complementary analysis, we also quantified global segregation using the system segregation index (SSI) (Chan et al., 2014) to assess whether the main pattern generalized to a standard graph-theoretical metric. It is further motivated by the necessity, for a RSN to be considered modular, to elicit increased processing while reducing its communication with other RSNs (Avena-Koenigsberger et al., 2018; Sporns & Betzel, 2016). Conversely, strong evidence for an integrated pattern was defined by the opposite configuration, reflecting the switch from internal processing to distributed processing. The presence of a positive within-network CB without reliably negative between-network CB was interpreted as only partial evidence of segregation, while the presence of a positive between-network CB without reliably positive within-network CB was interpreted as only partial evidence of integration.

Additionally, we explored whether each latent CB factor was supported by a network architecture differing from the intrinsic, RSN architecture described by the GINNA Atlas. Namely, separately for each latent CB factor, we performed hierarchical clustering on the 448 x 448 regional CB matrices (n=100 iterations) and extracted the clustering solution matching the resolution of the used RSN partition (n=34), resulting in the assignment of each region to one of 34 data-driven networks. Global estimates of network segregation and integration were then computed similarly to described above, using the data-driven network partitioning to define networks. The same post-hoc reliability analysis was applied to determine whether latent CB factors were associated with reliable network theoretical metrics effects across prediction repetitions (n=100) (95% confidence interval).

### 2.11 Statistical analysis

Statistical analyses related to specific analyses steps are detailed in the appropriate Methods sections.

## 3 Results

### 3.1 Functional connectivity prediction of behavior

First, we performed functional connectivity (FC) prediction of behavior to predict individual differences using each of the 58 behavioral measures. Using a half-split cross-validation procedure (Tian & Zalesky, 2021) together with a simple kernel ridge regression model (He et al., 2020), we were able to significantly predict 52 of the 58 included measures (paired t-tests, FDR q < 0.05) (Fig. 3, Table 1). Importantly, although 52 of the 58 behavioral measures were predicted significantly better than chance, the absolute predictive signal remained modest (sRMSE > 0.97). Thus, significance should be understood as indicating that FC-based predictions outperformed a permuted null model, rather than that they captured a large proportion of interindividual behavioral variance.

**Fig. 3.**
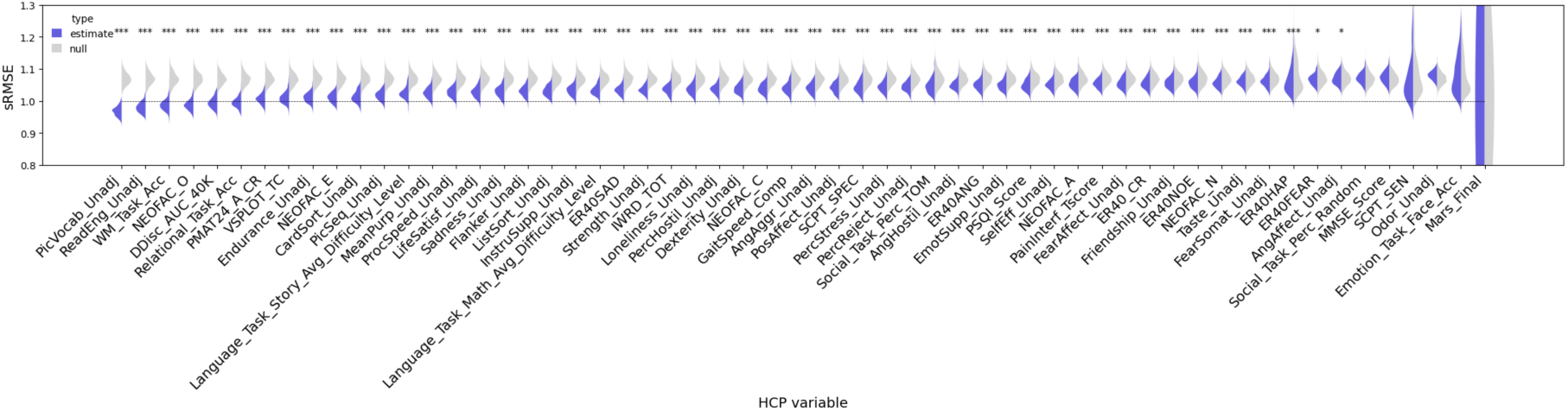
Prediction results across the 58 behavioral measures. Results are reported as standardized root mean squared error (sRMSE) after confounds regression performed on the prediction results (see *Methods*). Stars indicate significance, assessed by comparing the distribution of predictions (n=100) to a distribution of 100 permuted predictions (paired-t-tests) (Tian & Zalesky, 2021), with Benjamini-Hochberg false discovery rate correction for multiple comparisons (q < .05): ‘***’: q < .001; ‘**’; q < .01; ‘*’: q < .05.

**Table 1.** FC prediction of behavior performance.

| Behavioral measure | sRMSE | p | FDR q | Behavioral measure | sRMSE | p | FDR q |
| --- | --- | --- | --- | --- | --- | --- | --- |
| PicVocab_Unadj | 0.97 | <.001 | <.001 | AngAggr_Unadj | 1.04 | <.001 | <.001 |
| ReadEng_Unadj | 0.98 | <.001 | <.001 | PosAffect_Unadj | 1.04 | <.001 | <.001 |
| WM_Task_Acc | 0.99 | <.001 | <.001 | SCPT_SPEC | 1.04 | <.001 | <.001 |
| NEOFAC_O | 0.99 | <.001 | <.001 | PercStress_Unadj | 1.05 | <.001 | <.001 |
| DDisc_AUC_40K | 1.00 | <.001 | <.001 | PercReject_Unadj | 1.05 | <.001 | <.001 |
| Relational_Task_Acc | 1.00 | <.001 | <.001 | Social_Task_Perc_TOM | 1.05 | <.001 | <.001 |
| PMAT24_A_CR | 1.01 | <.001 | <.001 | AngHostil_Unadj | 1.05 | <.001 | <.001 |
| VSPLOT_TC | 1.01 | <.001 | <.001 | ER40ANG | 1.05 | <.001 | <.001 |
| Endurance_Unadj | 1.01 | <.001 | <.001 | EmotSupp_Unadj | 1.05 | <.001 | <.001 |
| NEOFAC_E | 1.01 | <.001 | <.001 | PSQI_Score | 1.05 | <.001 | <.001 |
| CardSort_Unadj | 1.01 | <.001 | <.001 | SelfEff_Unadj | 1.05 | <.001 | <.001 |
| PicSeq_Unadj | 1.02 | <.001 | <.001 | NEOFAC_A | 1.05 | <.001 | <.001 |
| Language_Task_Story_Avg_Difficulty_Level | 1.03 | <.001 | <.001 | PainInterf_Tscore | 1.06 | <.001 | <.001 |
| MeanPurp_Unadj | 1.03 | <.001 | <.001 | FearAffect_Unadj | 1.06 | <.001 | <.001 |
| ProcSpeed_Unadj | 1.03 | <.001 | <.001 | ER40_CR | 1.06 | <.001 | <.001 |
| LifeSatisf_Unadj | 1.03 | <.001 | <.001 | Friendship_Unadj | 1.06 | <.001 | <.001 |
| Sadness_Unadj | 1.03 | <.001 | <.001 | ER40NOE | 1.06 | <.001 | <.001 |
| Flanker_Unadj | 1.03 | <.001 | <.001 | NEOFAC_N | 1.06 | <.001 | <.001 |
| ListSort_Unadj | 1.03 | <.001 | <.001 | Taste_Unadj | 1.06 | <.001 | <.001 |
| InstruSupp_Unadj | 1.03 | <.001 | <.001 | FearSomat_Unadj | 1.06 | <.001 | <.001 |
| Language_Task_Math_Avg_Difficulty_Level | 1.04 | <.001 | <.001 | ER40HAP | 1.07 | <.001 | <.001 |
| ER40SAD | 1.04 | <.001 | <.001 | ER40FEAR | 1.07 | .018 | .020 |
| Strength_Unadj | 1.04 | <.001 | <.001 | AngAffect_Unadj | 1.07 | .028 | .031 |
| IWRD_TOT | 1.04 | <.001 | <.001 | Social_Task_Perc_Random | 1.07 | .584 | .616 |
| Loneliness_Unadj | 1.04 | <.001 | <.001 | MMSE_Score | 1.07 | 1.000 | 1.000 |
| PercHostil_Unadj | 1.04 | <.001 | <.001 | SCPT_SEN | 1.08 | .105 | .115 |
| Dexterity_Unadj | 1.04 | <.001 | <.001 | Odor_Unadj | 1.08 | 1.000 | 1.000 |
| NEOFAC_C | 1.04 | <.001 | <.001 | Emotion_Task_Face_Acc | 1.08 | .533 | .573 |
| GaitSpeed_Comp | 1.04 | <.001 | <.001 | Mars_Final | 2.53 | .740 | .766 |

Of note, prediction of Mars_Final performed particularly poorly (sRMSE = 2.53). This may reflect that contrast sensitivity is linked to fine-grained patterns within visual-sensitive regions (Yang et al., 2019), which may be better captured at a voxel-level resolution rather than by region-averaged connectivity measures. More generally, the non-significant measures may be weakly predicted because they capture processes with limited interindividual variability, strong state or sensory dependence, or neural signatures that are not well summarized by whole-brain region-averaged resting-state connectivity.

Overall, these findings indicate the presence of small but statistically reliable brain–behavior associations.

Similar results were obtained using the Schaefer2018 parcellation at 200 regions resolution (Schaefer et al., 2018), though with higher prediction error, as indicated by sRMSE values exceeding 1 (Supplementary Fig. 1).

### 3.2 Latent connectivity-behavior factors

To reveal the latent connectivity-behavior (CB) organization, Haufe matrices (i.e., connectivity features relevance for prediction) of significantly predicted behavioral measures were subsequently subjected to exploratory factor analysis (EFA). Parallel analysis revealed that the best solution for EFA computed on the 52 Haufe patterns was achieved for k = 10 dimensions (Fig. 4A). Although parallel analysis suggested up to 10 factors, inspection of the explained variance profile indicated a clear dominance of the first three factors, whereas several subsequent factors accounted for less than 5% of the common variance and contributed minimal incremental explanatory power (Fig. 4A). Consistent with recommendations emphasizing parsimony and substantive interpretability (Costello & Osborne, 2005; Fabrigar et al., 1999), these factors were not retained. The first three factors explained ∼25% of the variance (Fig. 4A), though this relatively modest proportion is expected given the weak and distributed nature of connectivity–behavior associations.

**Fig. 4.**
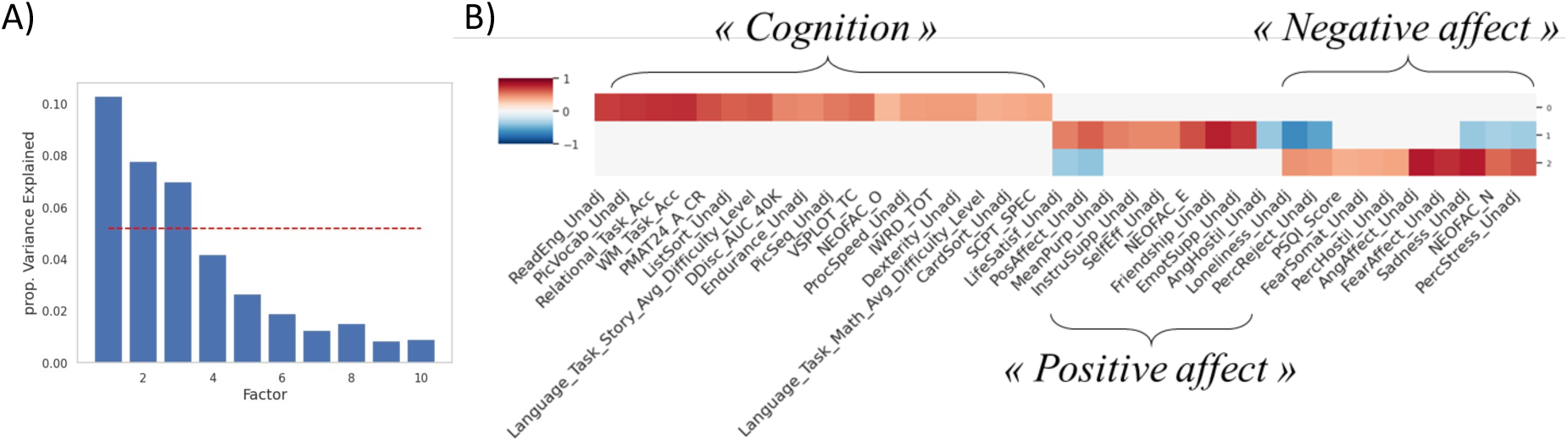
Latent connectivity-behavior factors. A) proportion explained variance for the exploratory factor analysis solution with 10 factors. Only the first three factors were retained (see *Methods*). B) loadings of each individual behavior Haufe matrices on the factors.

Replication of the analysis using the Schaefer200 parcellation atlas (Schaefer et al., 2018) resulted in a similar optimal solution of k = 10 factors, while only 2 factors explained a substantial share of the variance (> 5%) (Supplementary Fig. 2A). This is explained by the behavioral variables related to positive affect being regrouped with those related to negative affect in a single, bidirectional *Affect* factor, suggesting substantial anticorrelations in the FC patterns underlying positive and negative affect.

The three remaining latent CB factors (Fig. 4B) corresponded to a separation between:

#### Factor “*Cognition*”

Behavioral measures (n = 18 with |loadings| > 0.3, all positive) related to cognition (e.g., working memory task accuracy, picture vocabulary).

#### Factor “Positive affect”

Behavioral measures (n = 14, 8 positive) related to socio-emotional constructs, with strong positive loadings for variables with positive emotional valence with a social component (e.g., friendship, extraversion, emotional support) and negative loadings for variables with negative emotional valence (loneliness, perceived reject).

#### Factor “Negative Affect”

Behavioral measures (n = 12, 10 positive) referring to the experience of negative emotions (e.g., sadness, loneliness), opposed to variables with positive emotional valence (life satisfaction, positive affect).

### 3.3 Connectivity patterns predictive of cognition reflect global network segregation at rest

We first analyzed pooled estimates of network segregation and integration, i.e., averaged across all RSNs present in the GINNA atlas. Only the *Cognition* CB factor was associated with strong segregation (both a reliable positive effect for segregation and a negative effect for integration) at the global level (segregation: mean = 0.0127, 95%CI [0.0049, 0.0216]; integration: mean = −0.0015, 95%CI [-0.0025, −0.0007] (Fig. 5A, Table 2). No reliable effect for segregation or integration were found for *Positive Affect* (segregation: mean = 0.0018, 95%CI [-0.0087, 0.0120]; integration: mean = −0.0006, 95%CI [-0.0015, 0.0004]) or *Negative Affect* (segregation: mean = −0.0021, 95%CI [-0.0141, 0.0101]; integration: mean = 0.0006, 95%CI [-0.0005, 0.0016]) (Fig. 5A, Table 2).

**Fig. 5.**
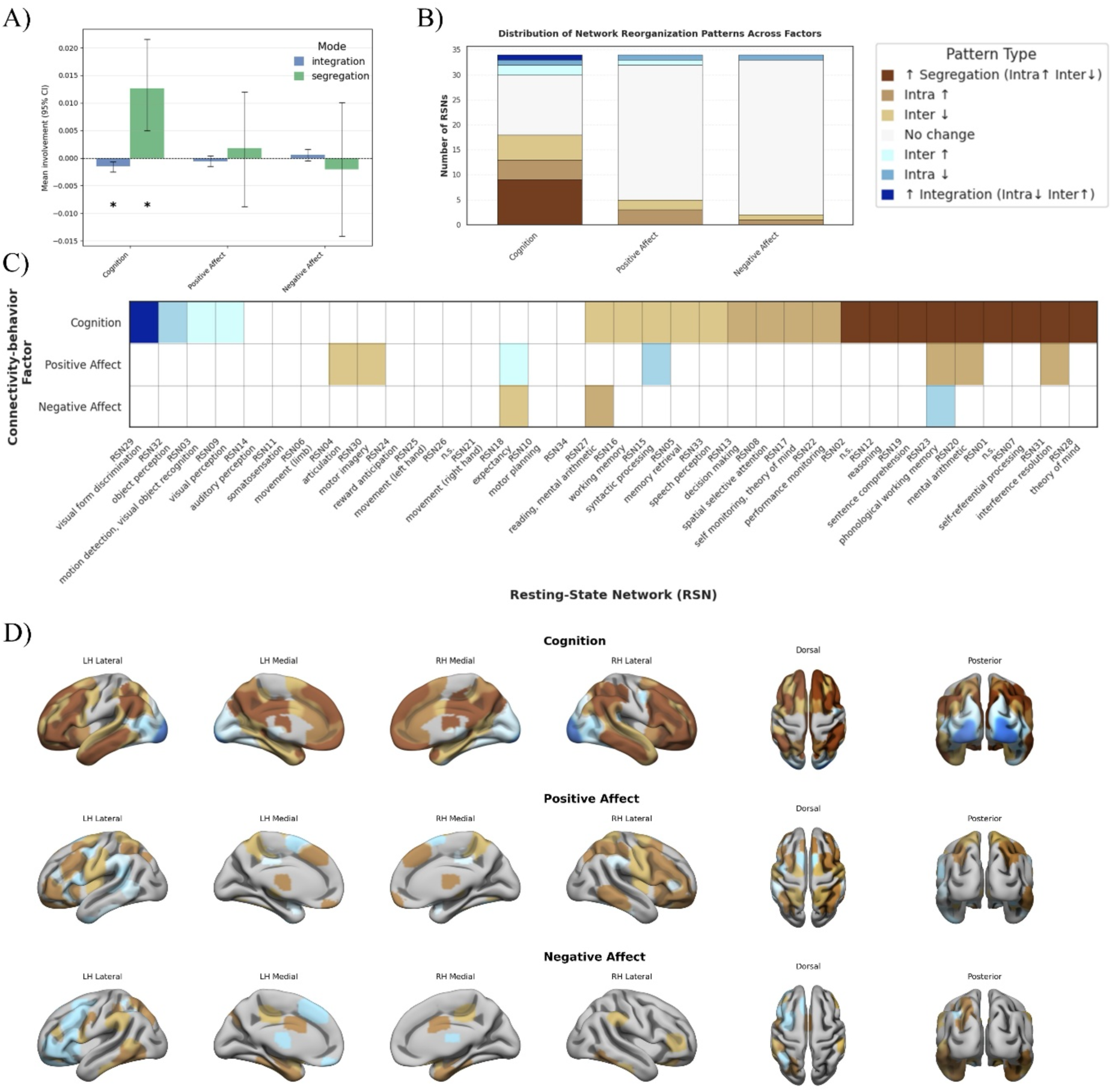
Network integration and network segregation across latent factors. A) Global network integration and segregation for each connectivity-behavior factor. B) Summary (cumulative count) of observed changes in network metrics at the RSN level. C) Changes in network metrics across the GINNA RSNs. D) surface projection of the observed changes in network metrics for each connectivity-behavior factor.

**Table 2.** Pooled network metrics for each latent connectivity-behavior factor.

| Latent CB factor | Network metric | mean | CI95% lower | CI95% upper | CI excludes 0 |
| --- | --- | --- | --- | --- | --- |
| Cognition | integration | -0.0015 | -0.0025 | -0.0007 | * |
|  | segregation | 0.0127 | 0.0049 | 0.0216 | * |
| Positive Affect | integration | -0.0006 | -0.0015 | 0.0004 |  |
|  | segregation | 0.0018 | -0.0087 | 0.0120 |  |
| Negative Affect | integration | 0.0006 | -0.0005 | 0.0016 |  |
|  | segregation | -0.0021 | -0.0141 | 0.0101 |  |

Replication using the Schaefer200 parcellation atlas (Schaefer et al., 2018) indicated that the segregation effect found for the *Cognition* CB factor was replicated at the Yeo 17 networks resolution (Yeo et al., 2011), but not at the 7 networks resolution, for which there was a reliable positive effect for segregation, but no reliable effect for integration (Supplementary Fig. 3).

As a complementary analysis, we repeated the global network analysis using the system segregation index (SSI). The Cognition factor was associated with a reliable positive SSI, whereas the Positive Affect and Negative Affect factors were associated with reliable negative SSI (Supplementary Fig. 4). Though this metric does not evidence the respective contributions of within vs between-network connectivity (i.e., a negative SSI can be caused by lower within-network connectivity, higher between-network connectivity, or a combination of both), these complementary results indicate that that the main findings were not primarily driven by the measures of segregation and integration used in the main analyses.

Complementary analyses revealed that replacing the RSN network partitioning by a data-driven network partitioning (k=34) of each latent CB matrices resulted in similar results. For the *Cognition* factor, better performance was associated with a reliable increase in global segregation (mean = 0.016, 95%CI [0.002, 0.031]) and a decrease in global integration (mean = −0.002, 95%CI [−0.004, −0.001]). For affective factors, similarly to the RSN partitioning, the data-driven network partitioning did not reveal any reliable effect for global segregation or integration, both for Positive Affect (segregation: mean = 0.007, 95%CI [−0.010, 0.022]; integration: mean = −0.0004, 95%CI [−0.002, 0.001]) and Negative Affect (segregation: mean = −0.004, 95%CI [−0.020, 0.012]; integration: mean = 0.001, 95%CI [−0.001, 0.002]) (Supplementary Fig. 1).

### 3.4 Connectivity patterns predictive of cognition reflect segregation of higher-level networks and integration of lower-level visual networks

In order to gain insights into the different involvement of RSNs for the latent CB factors, we investigated whether individual RSNs were associated with reliable changes in network integration or segregation. Overall, the *Cognition* CB factor was associated with the most effects in segregation and integration (n = 22 RSNs), while only few effects were uncovered for *Positive Affect* (n = 7) or *Negative Affect* (n = 3) (Fig. 5B).

We found that the *Cognition* CB factor was associated with strong segregation (i.e., both a reliable positive effect for segregation and a reliable negative effect for integration) for 9 RSNs (Fig. 5B). 7 of them were previously associated with tasks involving higher-level cognitive processes (Gillig et al., 2025): RSN07, associated with self-referential processing and corresponding to the canonical definition of the default mode network, RSN12 (reasoning), RSN19 (sentence comprehension), RSN20 (mental arithmetic), RSN23 (phonological working memory), RSN28 (theory of mind), and RSN31 (interference resolution) (Fig. 5C-D, Supplementary File 1). The two remaining ones were RSNs showing no significant association with task studies, namely RSN01, a dorsal subcomponent of the Default Mode Network, and RSN02, a distributed fronto-temporo-parietal network resembling a right-lateralized fraction of the domain-general network.

In addition, the *Cognition* CB factor was associated with weak segregation (positive effect for segregation without reliable effects for integration) for 4 RSNs (Fig. 5B): RSN08 (spatial selective attention), RSN13 (decision making), RSN17 (self-monitoring, theory of mind) and RSN22 (performance monitoring), as well as decreased integration for 4 RSNs: RSN05 (memory retrieval), RSN15 (syntactic processing), RSN16 (working memory), and RSN33 (speech perception) (Fig. 5C-D, Supplementary File 1).

Last, connectivity predictive of *Cognition* was associated with strong integration for RSN29 (visual form discrimination) and weak integration (positive effect for integration without reliable effects for segregation) for RSN03 (motion detection, visual object recognition), RSN09 (visual perception) and RSN32 (object perception) (Fig. 5C-D, Supplementary File 1).

### 3.5 Connectivity patterns predictive of affect do not show strong RSN-level segregation or integration

The *Positive Affect* CB factor was associated with weak segregation for two RSNs (Fig. 5B): RSN23 (phonological working memory) and RSN31 (interference resolution), a negative effect for integration for RSN04 (articulation) and RSN30 (motor imagery), as well as a negative effect for segregation for RSN18 (expectancy). Weak integration was evidenced for RSN15 (syntactic processing) (Fig. 5C-D, Supplementary File 1).

The *Negative Affect* CB factor was associated with weak segregation for RSN27 (reading, mental arithmetic), decreased segregation for RSN23 (phonological working memory), and a negative effect for integration for RSN18 (expectancy) (Fig. 5C-D, Supplementary File 1).

## 4 Discussion

Understanding how the intrinsic brain organization supports interindividual differences in behavior remains a central challenge in systems neuroscience. Using a combination of FC prediction of behavior, interpretability tools and network theoretical metrics, we provide evidence of the correspondence between the intrinsic, RSN brain architecture, and the FC architecture supporting behavior. Using a comprehensive set of behavioral measures from a large sample of healthy young adults, we show that behaviorally relevant connectivity patterns at rest organize into a small number of latent dimensions, and that these dimensions are supported by distinct large-scale architectural principles. Specifically, we show that behaviorally relevant FC organizes into 3 principal latent dimensions summarizing *Cognition*, *Positive Affect* and *Negative Affect*. Crucially, higher Cognition scores were associated with predictive connectivity patterns reflecting strong global resting-state network segregation, consistent with the interpretation that the intrinsic brain architecture may reflect a functionally specialized cognitive organization. By contrast, *Affect* did not and was instead associated with a negative system segregation index, suggesting that it is supported by a more flexible, distributed organization in which RSNs do not represent core affective modules. We furthermore highlight differential RSN involvements in *Cognition*, with strong segregation of RSNs associated with higher-level cognitive processes, and integration of lower-level, visual RSNs, suggesting a hierarchical organization of the intrinsic functional architecture that scaffolds cognitive performance.

### 4.1 Latent connectivity–behavior factors capture shared neural substrates across behaviors

The emergence of three latent connectivity–behavior factors indicates that predictive FC patterns are structured around a limited number of shared neural mechanisms despite the wealth of behavioral measures significantly predicted. Importantly, these factors were derived from connectivity–behavior mappings rather than behavioral measures alone, implying that they reflect common neural substrates underlying behavioral variance. The three latent factors combined to explain approximately 25% of the variance, reflecting the high dimensionality and heterogeneity of the connectivity-behavior space, which is inherently noisy due to the modest prediction performance observed. The *Cognition* factor aggregated diverse cognitive measures spanning working memory, reasoning, language, and processing speed, among others, consistent with the presence of shared neural components supporting cognitive performance (Chen et al., 2022). *Positive Affect* determinants were more associated with social-emotional components (extraversion, friendship, emotional support), while *Negative Affect* ones reflected more the experience of negative emotions (neuroticism, feelings of fear anger, and sadness), indicating that these dimensions are not exactly anticorrelated, but reflect distinct processes. However, replication using the Schaefer200 parcellation (Schaefer et al., 2018) resulted in these two affective factors to be regrouped into a single bidirectional *Affect* factor, suggesting that FC patterns underlying *Positive Affect* and *Negative Affect* remain substantially anticorrelated, or that this finding is sensitive to the parcellation scheme employed.

Notably, no factor corresponding uniquely to personality traits emerged, despite significant prediction of individual personality measures. Extraversion was included in the *Positive Affect* CB factor, Neuroticism in the *Negative Affect* factor, while Openness was included in *Cognition*. This shows that, within the present behavioral battery, personality-related connectivity patterns largely overlap with affective or cognitive dimensions rather than constituting their own latent axis. This contrasts with previous reports made on adolescents indicating shared common connectivity features for personality that largely differed from cognition (Chen et al., 2022).

### 4.2 Connectivity patterns predictive of cognition reflect global network segregation at rest

A central finding of this study is that the connectivity patterns associated with higher cognitive ability reflected greater global network segregation and reduced integration at rest. Importantly, networks were defined intrinsically (RSNs), not from their behavioral relevance. Additionally, this pattern remained consistent when the architecture was quantified using the system segregation index, indicating that the main result was not specific to our initial operationalization of segregation and integration. Together with the complementary analyses showing robustness under an alternative parcellation and network scheme at 17 Networks resolution, but not 7 Networks resolution (Schaefer et al., 2018; Yeo et al., 2011), this supports the robustness of the finding while highlighting the relevance of RSN granularity. Network segregation is widely thought to reflect functional specialization and efficient within-system processing (Avena-Koenigsberger et al., 2018; Benso et al., 2025; Sporns & Betzel, 2016), whereas integration dynamically supports coordination across systems when task demands require it (Bertolero et al., 2015; Cohen & D’Esposito, 2016; Shine et al., 2016; Shine & Poldrack, 2018; X. Wang et al., 2024). Within this framework, our results further suggest that connectivity patterns predictive of cognition are more consistent with a segregated intrinsic organization at rest, and therefore with the interpretation that RSNs may capture functionally specialized cognitive components (Barrett & Kurzban, 2006; Benso et al., 2025; Bertolero et al., 2015, 2018; Sporns & Betzel, 2016).

These findings do not contradict task-based evidence showing that complex cognitive performance requires increased integration across networks (Cohen & D’Esposito, 2016; Shine & Poldrack, 2018; X. Wang et al., 2024). Rather, they support a complementary account in which segregation and integration play distinct roles depending on brain state. While task execution necessitates transient integration to enable flexible coordination (Shine et al., 2016; Shine & Poldrack, 2018), the resting-state organization may reflect the long-term maintenance of a specialized architecture that reduces the need for large-scale reconfiguration during task engagement (Schultz & Cole, 2016). Consistent with this view, learning studies have shown that faster stimulus–response learning is associated with increased segregation at later learning stages, whereas earlier, goal-directed learning involves greater integration (Wang et al., 2024). Together, these findings suggest that large-scale changes in segregation at rest may index the offline consolidation or efficiency of overlearned processes, while integration remains a key mechanism to enable flexible adaptation to changing environmental demands.

### 4.3 Differential involvement of higher-level cognitive versus visual RSNs in *Cognition*

We further observed that the relationship between cognitive ability and network organization varied across RSNs. Specifically, higher cognitive ability was associated with predictive connectivity patterns reflecting strong segregation of multiple higher-order association networks implicated in self-referential processing, reasoning, language, working memory, theory of mind, and cognitive control (Gillig et al., 2025). This finding extends our current as well as prior evidence for shared neural substrates across *Cognition* (Chen et al., 2022) by showing that cognitive ability is associated with connectivity patterns reflecting coordinated segregation across a distributed set of higher-level RSNs, rather than by a single network or localized mechanism.

Importantly, the consistency of segregation effects across multiple RSNs within a latent cognition factor suggests that cognitive performance relies on shared but non-specific architectural principles across networks. This aligns with evidence that RSNs do not map one-to-one onto task-defined functional systems (Davis et al., 2017; Thompson & Fransson, 2017). Instead, our results support the view that RSNs are not strictly encapsulated cognitive modules (Benso et al., 2025; Noble et al., 2024), but instead represent more elementary processing units, that are flexibly combined during task execution. Supporting this view, task-evoked activation patterns are partly predicted by common resting-state components (Scholz et al., 2024), and, although better task performance is associated with reduced reconfiguration between rest and task, substantial reorganization remains necessary to meet task demands (Davis et al., 2017; Salehi et al., 2020; Schultz & Cole, 2016).

In contrast, predictive connectivity patterns linked to higher cognitive ability reflected greater integration of several visual networks. This pattern suggests that lower-level visual systems may play a more integrative role within the intrinsic architecture associated with cognition, potentially facilitating efficient information transfer to higher-order systems. Such an interpretation is consistent with evidence that resting-state connectivity constrains task-evoked information flow and supports activity propagation during cognitive processing (Ito et al., 2017). Together, these results suggest that intrinsic RSN organization may provide a hierarchical scaffold that supports cognitive performance through differentiated roles of segregation and integration.

### 4.4 Absence of strong segregation signatures for affective factors

Unlike cognition, affective connectivity-behavior dimensions were not associated with reliable global effects in network segregation or integration, and only partial, network-specific effects were observed. Because GINNA’s intrinsic RSN structure has been shown to relate to known cognitive components (Gillig et al., 2025), a possibility remained that the RSN composition used to compute segregation and integration metrics may be optimized for the cognitive dimension, and that different networks may underlie the affective dimension. To test this possibility, new network compositions were obtained by a data-driven clustering independently for each dimension. Interestingly, this change did not alter the results, and no new segregation or integration effects were uncovered. This suggests that the lack of network segregation for the predictive connectivity associated with *Affect* does not only relate only to a misalignment with the intrinsic network architecture, but rather, that the FC basis of *Affect* is characterized by an absence of homogeneity within and between-systems (*i.e.*, clusters regroup both positive and negative edges). All in all, while the present evidence suggest that the intrinsic organization into RSNs provides a basis for a modular cognitive organization, it does not support RSNs as core affective modules (Dan et al., 2023; Lindquist et al., 2012; Lindquist & Barrett, 2012). This aligns with contemporary views that emotional processes are not supported by specific RSNs, but by a reconfiguration of whole-brain, distributed functional networks that overlap with systems involved in processes including, but not limited to, conceptualization, attention, motor function, and executive control (Dan et al., 2023; Lindquist et al., 2012; Lindquist & Barrett, 2012; Pessoa, 2017).

Another possibility is that affective traits rely on more distributed and context-sensitive interactions that are not well captured by static segregation metrics. Alternatively, affective variance may be more state-dependent, reducing the stability of trait-level intrinsic signatures. More generally, these findings are consistent with evidence that resting-state functional connectivity has limited or variable predictive power for certain individual differences, particularly outside the cognitive domain (Chen et al., 2022; Ooi et al., 2022; Sui et al., 2020; Wu et al., 2023). While this hinders our ability to firmly conclude on the absence of an intrinsic segregated architecture associated with *Affect*, our complementary findings of a reliable negative system segregation index for *Affect* suggests that the weak connectivity features predicting affective measures primarily distribute outside of RSNs as defined in the present study. Though network segregation was also not observed using a data-driven partitioning, this does not rule out the alternative possibility that a segregated pattern might appear with sufficient predictive power. Additionally, the absence of strong segregation effects does not necessarily imply a lack of neural organization for affect, and rather, the present findings suggest that its architecture may not be primarily driven by the intrinsic RSN architecture, or by their large-scale segregation–integration balance.

### 4.5 Limitations and methodological considerations

Several limitations warrant consideration. First, the present results are necessarily limited by the modest predictive strength of the models. Although many behavioral measures were predicted significantly better than chance, the absolute amount of explained variance remained small, meaning that the connectivity–behavior associations identified here reflect a small but statistically reliable signal rather than strong predictive accuracy. As in all FC prediction of behavior studies, this high prediction error directly constrains model interpretability, since the reliability and biological meaning of feature weights depend on prediction performance (Chen et al., 2023; Wu et al., 2023). This limitation propagates across the successive steps of our pipeline, from prediction to Haufe inversion, latent factor analysis, and network-level summaries. Nonetheless, our conclusions are based on out-of-sample testing combined with a reliability analysis of feature weights across repetitions of the prediction, which partially, but not completely, mitigates this issue. Accordingly, the present findings should be interpreted cautiously, as describing broad organizational tendencies in behaviorally relevant connectivity patterns rather than providing strong or mechanistic evidence about how intrinsic brain architecture determines behavior. Additionally, the kernel regression model, chosen for its state-of-the-art performance (He et al., 2020; Schulz et al., 2020), operates on the space of global similarities between whole-brain FC matrices, therefore reducing the ability to detect sparse, local effects.

Second, because latent connectivity–behavior factors depend on the behavioral space sampled, the present findings should be interpreted with respect to this specific 58-measure panel. In particular, differences between cognition and affect may partly reflect differences in construct heterogeneity, psychometric reliability, and sensitivity of the selected measures, in addition to differences in intrinsic functional organization. Next, we used a discrete RSN partition which highlights modular specialization, but may underrepresent hub regions, despite their known relevance for cognition (Bertolero et al., 2018) and in the underlying dynamics of the resting state (Saggar et al., 2022). Gradient-based approaches better capture these intermediary positions and continuous transitions (Margulies et al., 2016). Our findings therefore should not be taken to imply that intrinsic brain organization is exclusively discrete, and future work should test whether this cognitive effect is complemented by hub- or gradient-based signatures of large-scale integration.

An additional limitation concerns residual head motion. Because motion can increase short-range connectivity while decreasing longer-range connectivity, it could bias within-versus between-network summaries and thus affect segregation-related metrics (Power et al., 2012; Satterthwaite et al., 2012). Although motion was addressed through nuisance regression, stringent exclusion, and frame censoring prior to FC estimation, residual effects may remain. Recent work suggests that, after stringent censoring, such effects may more often attenuate or distort than spuriously inflate brain-behavior associations, but do not disappear entirely (Kay et al., 2025; Parkes et al., 2018; Pavlovich et al., 2025). Given that prediction performance for Affect was lower than for Cognition, the absence of reliable segregation effects for Affect may partly reflect reduced signal-to-noise in the underlying brain–behavior associations, which could be more susceptible to residual motion-related biases.

Last, static resting-state FC cannot capture dynamic reconfiguration or causal interactions. Future work integrating task-based and dynamic connectivity analyses (Cutts et al., 2025), as well as longitudinal and interventional approaches, will be critical for clarifying how intrinsic segregation relates to cognition across brain states.

## 5 Data and code availability

Data were provided by the Human Connectome Project, WU-Minn Consortium (Principal Investigators: David Van Essen and Kamil Ugurbil; 1U54MH091657) funded by the 16 NIH Institutes and Centers that support the NIH Blueprint for Neuroscience Research; and by the McDonnell Center for Systems Neuroscience at Washington University.

Code used for analyses is openly available at https://github.com/Achillegillig/rsn-segregation-behavior.

## 6 Author Contributions

Conceptualization: A.G., M.J.; Methodology: A.G., M.J., G.J., S.C.; Formal analysis: A.G.; Visualization: A.G., M.J.; Writing—original draft: A.G.; Writing—review & editing: G.J., M.J., SC.

## Acknowledgements

Computer time for this study was provided by the computing facilities of the MCIA (Mésocentre de Calcul Intensif Aquitaine, Bordeaux, France).

