## Supplementary Table1, Supplementary File 1, Supplementary Fig. 1, Supplementary Fig. 2A, Supplementary Fig. 3, Supplementary Fig. 4 for "Connectivity patterns predictive of cognition, but not affect, reflect a segregated intrinsic network architecture"

| HCP Variable | Extended name | HCP Variable | Extended name |
|---|---|---|---|
| PicSeq_Unadj | Visual Episodic Memory | WM_Task_Acc | Working Memory (N-back) |
| CardSort_Unadj | Cognitive Flexibility | NEOFAC_A | Agreeableness (NEO) |
| Flanker_Unadj | Inhibition (Flanker Task) | NEOFAC_O | Openness (NEO) |
| PMAT24_A_CR | Fluid Intelligence | NEOFAC_C | Conscientiousness (NEO) |
| ReadEng_Unadj | Vocabulary (Pronunciation) | NEOFAC_N | Neuroticism (NEO) |
| PicVocab_Unadj | Vocabulary (Picture Matching) | NEOFAC_E | Extroversion (NEO) |
| ProcSpeed_Unadj | Processing Speed | ER40_CR | EmotionRecog. - Total |
| DDisc_AUC_40K | Delay Discounting | ER40ANG | EmotionRecog. - Anger |
| VSPLOT_TC | Spatial Orientation | ER40FEAR | EmotionRecog. - Fear |
| SCPT_SEN | Sustained Attention - Sens. | ER40HAP | EmotionRecog. - Happiness |
| SCPT_SPEC | Sustained Attention - Spec. | ER40NOE | EmotionRecog. - Neutral |
| IWRD_TOT | Verbal Episodic Memory | ER40SAD | EmotionRecog. - Sadness |
| ListSort_Unadj | Working Memory (List Sorting) | AngAffect_Unadj | Anger - Affect |
| MMSE_Score | Cognitive Status (MMSE) | AngHostil_Unadj | Anger - Hostility |
| PSQI_Score | Sleep Quality | AngAggr_Unadj | Anger - Aggressiveness |
| Endurance_Unadj | Walking Endurance | FearAffect_Unadj | Fear - Affect |
| GaitSpeed_Comp | Walking Speed | FearSomat_Unadj | Fear - Somatic Arousal |
| Dexterity_Unadj | Dexterity | Sadness_Unadj | Sadness |
| Strength_Unadj | Grip Strength | LifeSatisf_Unadj | Life Satisfaction |
| Odor_Unadj | Odor Identification | MeanPurp_Unadj | Meaning of Life |
| PainInterf_Tscore | Pain Interference Survey | PosAffect_Unadj | Positive Affect |
| Taste_Unadj | Taste Intensity | Friendship_Unadj | Friendship |
| Mars_Final | Contrast Sensitivity | Loneliness_Unadj | Loneliness |
| Emotion_Task_Face_Acc | Emotion Face Matching | PercHostil_Unadj | Perceived Hostility |
| Language_Task_Math_Avg_Difficulty_Level | Arithmetic | PercReject_Unadj | Perceived Rejection |
| Language_Task_Story_Avg_Difficulty_Level | Story Comprehension | EmotSupp_Unadj | Emotional Support |
| Relational_Task_Acc | Relational Processing | InstruSupp_Unadj | Instrumental Support |
| Social_Task_Perc_Random | Social Cognition - Random | PercStress_Unadj | Perceived Stress |
| Social_Task_Perc_TOM | Social Cognition - Interaction | SelfEff_Unadj | Self-Efficacy |

**Supplementary Table 1 List of the 58 selected behavioral measures from the Human Connectome Project dataset.** We used the same selection, spanning cognitive, emotional and social-behavioral domains, as in (Liégeois et al., 2019)

| |
|---|
| https://www.dropbox.com/scl/fi/bndt3kurjwd71byvdb9ik/segregation_integration_table.xlsx?rlkey=cv4f49belya2ima9mpymywmvl&st=q2svuap8&dl=0 |
| **Supplementary File 1 network segregation / integration table for each RSN.** |

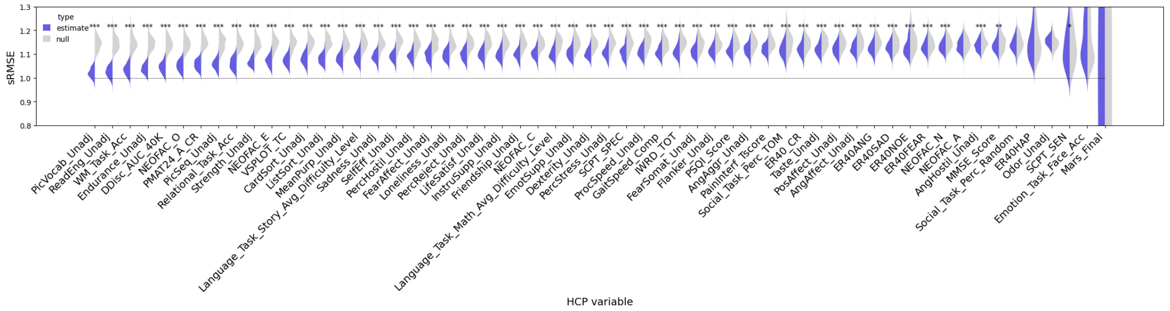


**Supplementary Fig. 1 Functional connectivity prediction of behavior replication using Schaefer 200 parcellation.** The figure indicates that the functional connectivity prediction of behavior procedure is stable under a different parcellation scheme, with only one variable (SCPE_SEN) becoming significant when it was originally not.

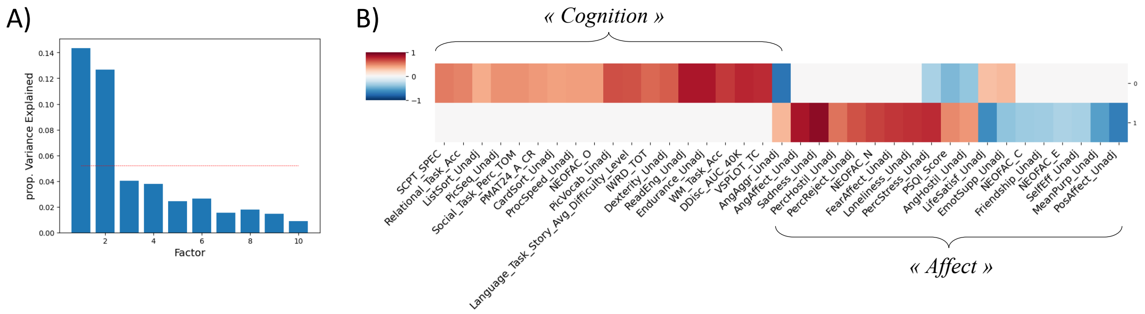


**Supplementary Fig. 2 Latent connectivity-behavior factor analysis replication using Schaefer 200 parcellation.** A) proportion explained variance for the exploratory factor analysis solution with 10 factors. Contrary to the solution obtained using the original parcellation atlas, two factors appear to be dominant. B) loadings of each individual behavior Haufe matrices on the factors. Replication using the Schaefer200 parcellation led the original "Positive Affect" and "Negative Affect" factors to be regrouped into a single component, with anticorrelated loadings between behavioral measures associated with positive emotional valence and those associated with negative emotional valence.

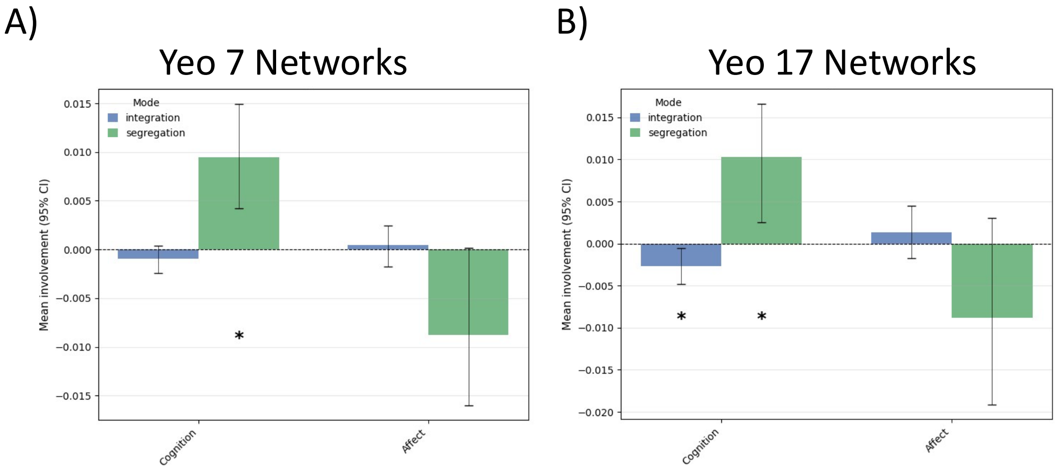


**Supplementary Fig. 3 Global network analysis replication using Yeo 7 / 17 Networks.** Replication reveals that strong segregation (increased segregation together with decreased segregation) for *Cognition* could be replicated using the Yeo 17 Networks resolution, but not the 7 Networks resolution.

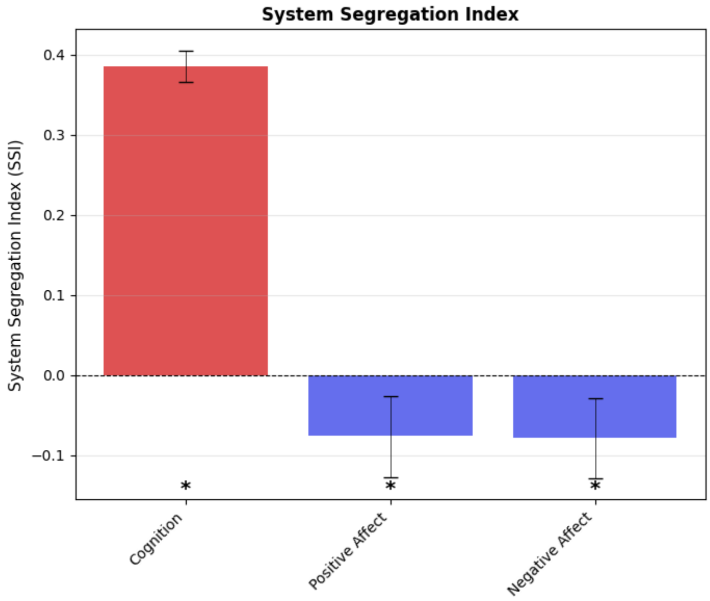


**Supplementary Fig. 4 Global network analysis using system segregation index (SSI) as a metric.** Results show that the *Cognition* factor is associated with a reliable positive SSI, indicating higher within-network relevance as compared to between-network connectivity. By contrast, *Positive Affect* and *Negative Affect* are associated with a reliable negative SSI, suggesting a distributed organization of connectivity relevant for *Affect*. Error bars indicate the 95% confidence interval. Stars indicate a reliable effect across prediction repetitions (95% confidence interval).

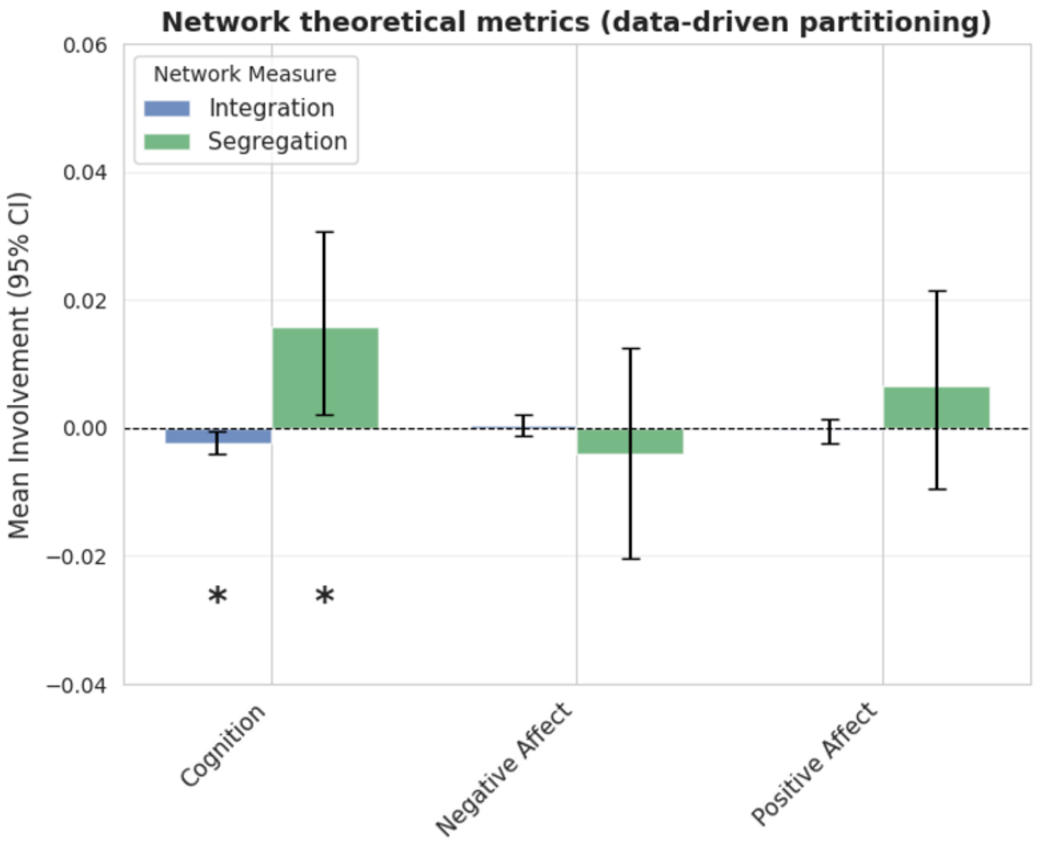


**Supplementary Fig. 4 Data-driven partitioning does not reveal global network segregation or integration for *Affect*.** The Fig. shows, for each latent CB factor, global changes in network integration and segregation for each connectivity-behavior factor across prediction repetitions (n=100) based on a data-driven network partitioning solution (k=34). Error bars indicate the 95% confidence interval. Stars indicate a reliable effect across prediction repetitions (95% confidence interval).
